# Processivity and BDNF-dependent modulation of signalling endosome axonal transport are impaired in aged mice

**DOI:** 10.1101/2025.01.30.635507

**Authors:** David Villarroel-Campos, Elena R. Rhymes, Andrew P. Tosolini, Bilal Malik, Alessio Vagnoni, Giampietro Schiavo, James N. Sleigh

**Author notes:** Equal contribution (David Villarroel-Campos, Elena R. Rhymes, Andrew P. Tosolini).

## Abstract

A healthy nervous system is reliant upon an efficient transport network to deliver essential cargoes throughout the extensive and polarised architecture of neurons. The trafficking of cargoes, such as organelles and proteins, is particularly challenging within the long projections of neurons, which, in the case of axons, can be more than four orders of magnitude longer than cell bodies. It is therefore unsurprising that disruptions in axonal transport have been reported across neurological diseases. A decline in this essential process has also been identified in many aging models, perhaps compounding age-related neurodegeneration. Via intravital imaging, we recently determined that, despite a reduction in overall motility, the run speed and displacement of anterograde mitochondrial transport were unexpectedly enhanced in aged mouse peripheral nerves. Here, to determine how aging impacts a different axonal cargo, we evaluated *in vivo* trafficking of signalling endosomes in motor axons of mouse sciatic nerves from 3 to 22 months. Contrasting with mitochondria, we did not detect alterations in signalling endosome speed, but found a consistent rise in pausing that manifested after 18 months. We then treated muscles with brain-derived neurotrophic factor (BDNF), which regulates axonal transport of signalling endosomes in motor neurons; however, we observed no change in the processivity defect at 22 months, consistent with downregulation of the BDNF receptor TrkB at the neuromuscular junction. Together, these findings indicate that aging negatively impacts signalling endosome trafficking in motor axons likely through dampened BDNF signalling at the motor neuron-muscle interface.

**Graphical Abstract:** 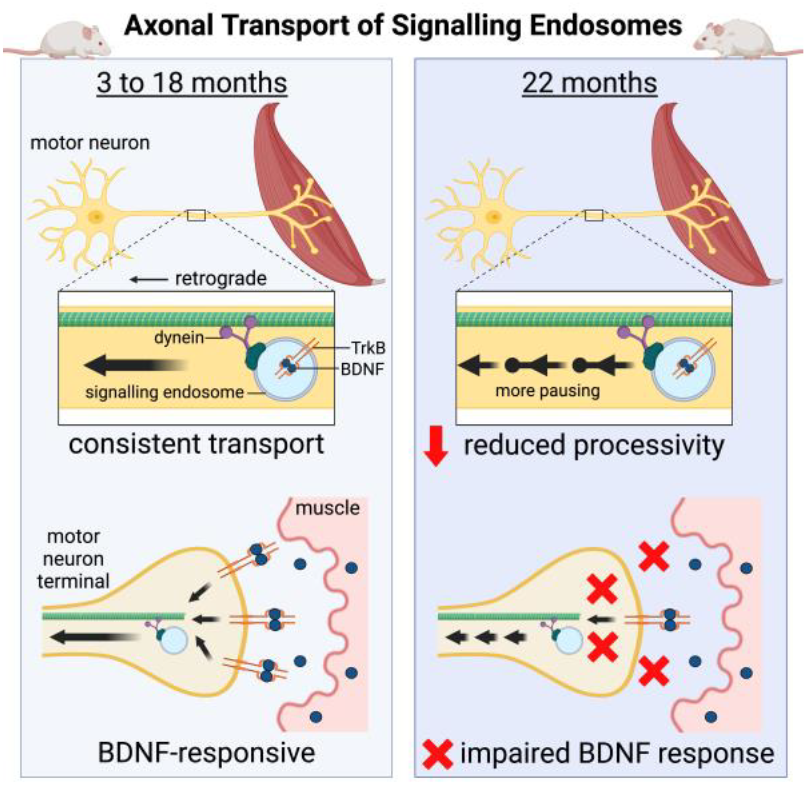

## Introduction

The highly specialised morphology of neurons comprises dendrites and axons, which are cytoplasmic projections that increase the cell surface area, enable the formation of complex neuronal networks, and expand the distance over which electrical signals can be transmitted. Axons can extend in excess of a metre; hence, neurons rely on a specialised transport system for delivery of diverse cargoes between cell bodies and axon terminals to maintain homeostasis and survival. This process, known as axonal transport, is dependent upon motor proteins, which selectively capture cargoes and actively engage the microtubule cytoskeleton for bi-directional trafficking (Maday et al., 2014). Anterograde transport connects the cell body to axon terminals and is reliant upon the kinesin family of motors (Hirokawa et al., 2009), whereas retrograde transport is oriented in the opposite direction and requires cytoplasmic dynein (Reck-Peterson et al., 2018). Axonal transport can be broadly divided into fast and slow categories: fast axonal transport delivers both membranous (e.g., mitochondria, secretory vesicles and signalling endosomes) and membrane-less organelles at a rate of ≈50-200 mm/day, whereas slow axonal transport conveys cytosolic/cytoskeletal proteins at the much slower rate of ≈0.2-10 mm/day due to prolonged and frequent pauses (Roy, 2020). Perturbations in axonal transport have been identified in many different neurological diseases, and mutations in the trafficking machinery can cause neurodegeneration, hence the biological importance of this process is well established (Brady and Morfini, 2017; Sleigh et al., 2019).

Aging is the primary risk factor for most neurodegenerative diseases, and some neuropathological signatures can mirror those underpinning aging (Hou et al., 2019). The impact of aging on axonal transport has been studied for many decades (Mattedi and Vagnoni, 2019), initially using methods such as nerve ligation and injection of radiolabelled tracers, and more recently via live imaging of fluorescent markers (Surana et al., 2020). Assessment of axonal transport *en masse* using classic techniques that rely on indirect biochemical analyses has repeatedly revealed a decline with age in both fast and slow axonal transport in central and peripheral nerves of rodents (Brunetti et al., 1987; Castel et al., 1994; Fernandez and Hodges-Savola, 1994; Frolkis et al., 1997; Frolkis et al., 1985; Geinisman et al., 1977; Goemaere-Vanneste et al., 1988; Hoffman et al., 1983; Komiya, 1980; McMartin and O’Connor, 1979; McQuarrie et al., 1989; Stromska and Ochs, 1982), which has been corroborated using computed tomography and magnetic resonance imaging approaches (Cross et al., 2008; Lee et al., 2022; Minoshima and Cross, 2008; Terry et al., 2022); however, these methods cannot differentiate between motor and sensory neurons and are impacted by several transport-independent features, including efficiency of tracer uptake and axon density.

Building on this early work, several studies using specific fluorescent cargoes have confirmed that, in some contexts, the processivity of axonal transport is perturbed as animals age. For instance, in *Caenorhabditis elegans*, an age-dependent, progressive reduction is observed in the speed of synaptic vesicle and mitochondrion trafficking within motor and sensory axons, respectively (Li et al., 2016; Morsci et al., 2016). Similarly, in mice, there is a decline between 18 and 24 months in the velocity of nicotinamide mononucleotide adenylyltransferase 2 (NMNAT2)-positive vesicles in sciatic nerve explants (Milde et al., 2015), whereas *in vivo* mitochondrion speed and distance of transport are diminished at 23-25 months in intact axons of retinal ganglion cells (Takihara et al., 2015). However, a plethora of publications indicate that not all cargoes or neuronal subtypes display age-related alterations in axonal velocities. For example, NMNAT2-positive vesicle speeds were unaltered in 24 month-old optic and hippocampal nerve explants (Milde et al., 2015), autophagic vesicle dynamics remain largely unchanged at 24 months in distal compartments of mouse sensory neurons *in vitro* (Tsong et al., 2023), and mitochondrial transport velocities remain unchanged across ages in sensory axons of the *Drosophila* wing (Vagnoni et al., 2016), mouse tibial nerve explants (Gilley et al., 2012), and mouse primary sensory neurons (Sleigh et al., 2024). Nevertheless, despite the lack of change in the processivity of axonal cargoes, these studies all showed a clear reduction in the overall number of transported cargoes, indicating that aging does not necessarily impact functionality of the trafficking machinery, but instead impairs processes that control the initiation or maintenance of axonal transport. Supporting this idea, the negative impact of aging on transport can be experimentally reversed, as shown by boosting cAMP/PKA signalling in *Drosophila* wing neurons (Vagnoni and Bullock, 2018) and in response to nerve injury in mouse tibial nerve explants (Milde et al., 2015).

The nature of reported axonal transport disruptions does not appear to follow an organelle- or neuron-specific pattern; thus, the impact of aging is context-dependent and requires systematic assessment across experimental systems to precisely dissect the relationship between age and axonal transport. It is also worth considering that removal of neurons from their natural milieu of interacting cells and molecules can alter transport dynamics, suggesting that *in vivo* approaches that enable transport assessment in situ provide a powerful platform for evaluating the effects of tissue and systemic aging (Sleigh et al., 2017).

Mitochondria are the most well studied axonal organelle (Saxton and Hollenbeck, 2012); however, they are bi-directionally trafficked, pause frequently and can remain stationary for long periods, hence their kinetic properties are dissimilar to other cargoes, such as intracellular vesicles, including signalling endosomes, which display highly processive retrograde movements and distinct regulatory mechanisms (Gibbs et al., 2015; Maday et al., 2014). We recently showed that mitochondria from mice aged 19-22 months display increased anterograde run speed and displacement within intact sciatic nerve axons compared to 3-4 month-old animals (Sleigh et al., 2024). Here, we made use of our intravital imaging approach to assess whether the processivity of axonal transport of a second organelle, the signalling endosome, is also altered with advanced age in mouse peripheral nerves.

## Materials and Methods

### Ethical Approval

Experiments involving mice were performed under license from the UK Home Office in accordance with the Animals (Scientific Procedures) Act (1986) and were approved by the UCL Queen Square Institute of Neurology Ethical Review Committee.

### Animals

Mice on the C57BL/6J background were maintained under a 12 h light/dark cycle at a constant room temperature of ≈21°C with water and food ad libitum (Teklad global 18% protein rodent diet, Envigo, 2018C). Cages were enriched with nesting material, plastic/cardboard tubes and wooden chew sticks. Males and females were assessed at 3 (90-115 days old), 9 (251-279 days old), 18 (555-615 days old) and 22 (663-687 days old) months of age (**Table S1**). Data from previously published studies were included within the presented figures, and are re-used under the CC BY 4.0 licence (https://creativecommons.org/licenses/by/4.0) (Rhymes et al., 2024; Sleigh et al., 2020a; Sleigh et al., 2023) (**Table S1**).

### In vivo axonal transport imaging

An atoxic binding fragment of the tetanus neurotoxin (H_C_T, residues 875-1,315 fused to a cysteine-rich tag and a human influenza haemagglutinin epitope) was used to perform live imaging of signalling endosome axonal transport (**Figure 1A**), as described in detail elsewhere (Gibbs et al., 2016; Sleigh et al., 2020b; Tosolini et al., 2021). Briefly, imaging was performed using either an LSM 780 or LSM 980 confocal microscope (Zeiss) equipped with a pre-warmed environmental chamber set to 38°C. Images were acquired every ≈3 seconds using a 63× Plan-Apochromat oil immersion objective (Zeiss). For experiments without brain-derived neurotrophic factor (BDNF), H_C_T was injected under isoflurane-induced anaesthesia into the tibialis anterior on one side of the body and the contralateral gastrocnemius; this enabled transport assessment in peripheral nerves supplying distinct hindlimb muscles of the same mouse, as performed previously (Rhymes et al., 2024; Sleigh et al., 2023). The side of injection (tibialis anterior versus gastrocnemius) was alternated between mice to eliminate time-under-anaesthesia and right/left biases. Imaging was performed 4-8 h later in both sciatic nerves. For muscle supplementation experiments, 25 ng recombinant human BDNF (Peprotech, 450-02) was pre-mixed and co-administered with H_C_T-555 into both the gastrocnemius and tibialis anterior muscles of the right leg, before imaging the right sciatic nerve 4-8 h later. Thicker H_C_T-positive axons were selected to prioritise evaluation of transport in motor axons (Sleigh et al., 2020a). Axonal transport at the node of Ranvier is slowed (Tosolini et al., 2024), thus these structures were avoided. Imaging of transport was completed within 2 h of inducing terminal anaesthesia.

**Figure 1.**
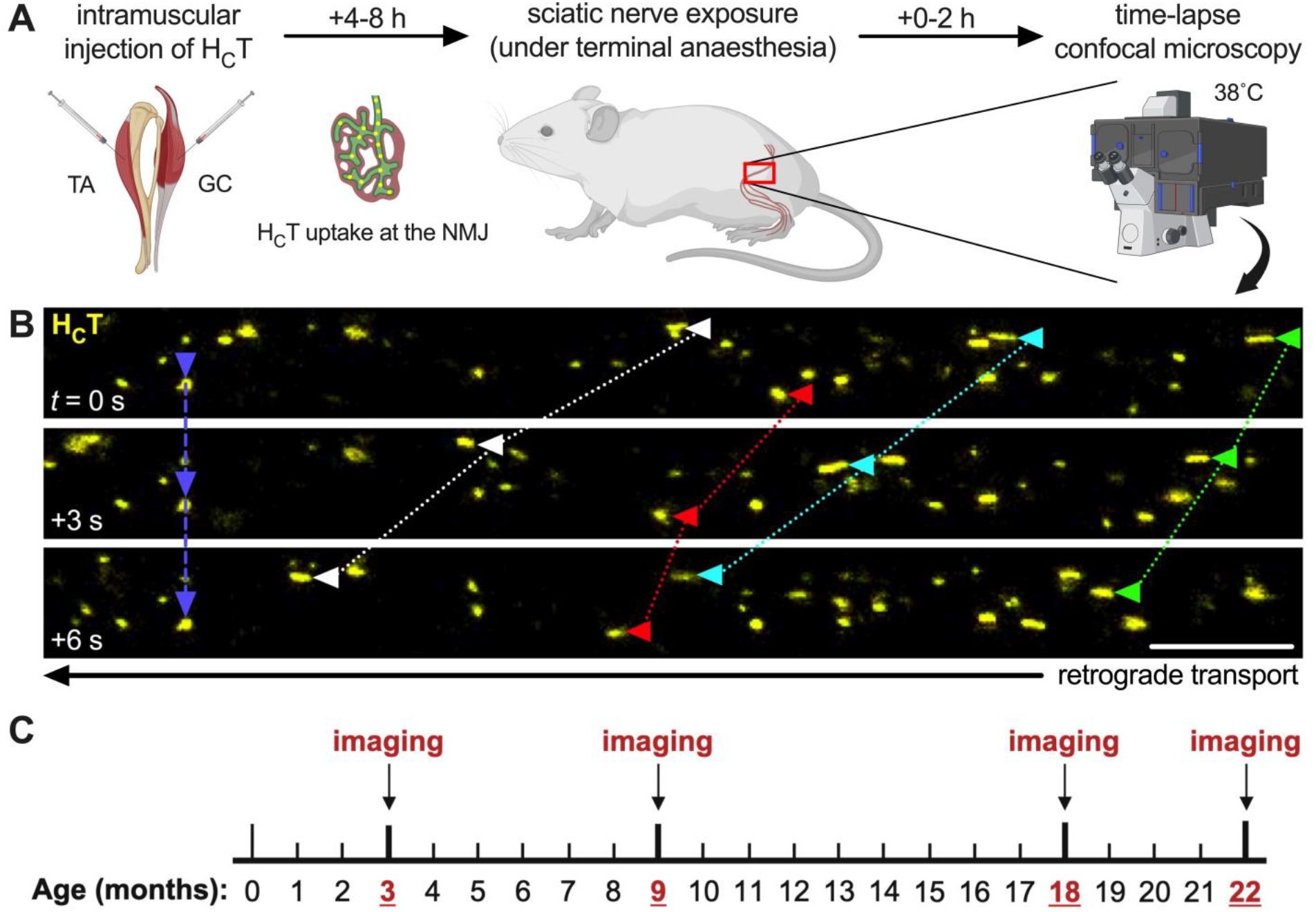
Imaging *in vivo* axonal transport of H_C_T-positive signalling endosomes in mice aged 3 to 22 months. (**A**) Upon intramuscular injection, fluorescently labelled H_C_T is taken up at the neuromuscular junction (NMJ) prior to retrograde axonal transport within signalling endosomes, which can be visualised 4-8 h later in intact peripheral axons of exposed sciatic nerves using time-lapse confocal microscopy. *GC*, gastrocnemius; *TA*, tibialis anterior. (**B**) Retrogradely transported H_C_T-positive endosomes (pseudo-coloured yellow) are individually tracked to quantitatively assess their dynamics. Colour-coded arrowheads identify five individual endosomes, one of which remains stationary across the three frames. (**C**) Timeline of study – *in vivo* imaging was performed at 3, 9, 18 and 22 months of age with and without muscle supplementation with BDNF.

### In vivo axonal transport analysis

A minimum of 10 endosomes per axon and at least three axons per mouse were manually tracked using the TrackMate plugin on ImageJ (https://imagej.net/ij/, **Figure 1B**) (Tinevez et al., 2017). Only endosomes that could be tracked for ≥5 consecutive frames were analysed – cargoes that did not fit this criterion were omitted due to going out of focus or being obscured by breathing artefact. An endosome was determined to have paused if it remained stationary (speed < 0.1 μm/sec) for two consecutive frames (3 sec). These pauses are included in the calculated speeds for each endosome. Endosomes that paused for ≥10 consecutive frames (≈ 27 sec) were rare (<1%) and excluded from the analyses, as the majority do not regain motility during the imaging period. Endosomes that moved solely anterogradely were also rare (<1%) and excluded, as we are interested in the retrograde dynamic properties of these cargoes. The “% time paused” is a calculation of the time all tracked endosomes remained stationary. To generate the combined tibialis anterior and gastrocnemius analyses presented in Figure 2J-L, the data from the two separately injected muscles were pooled.

**Figure 2.**
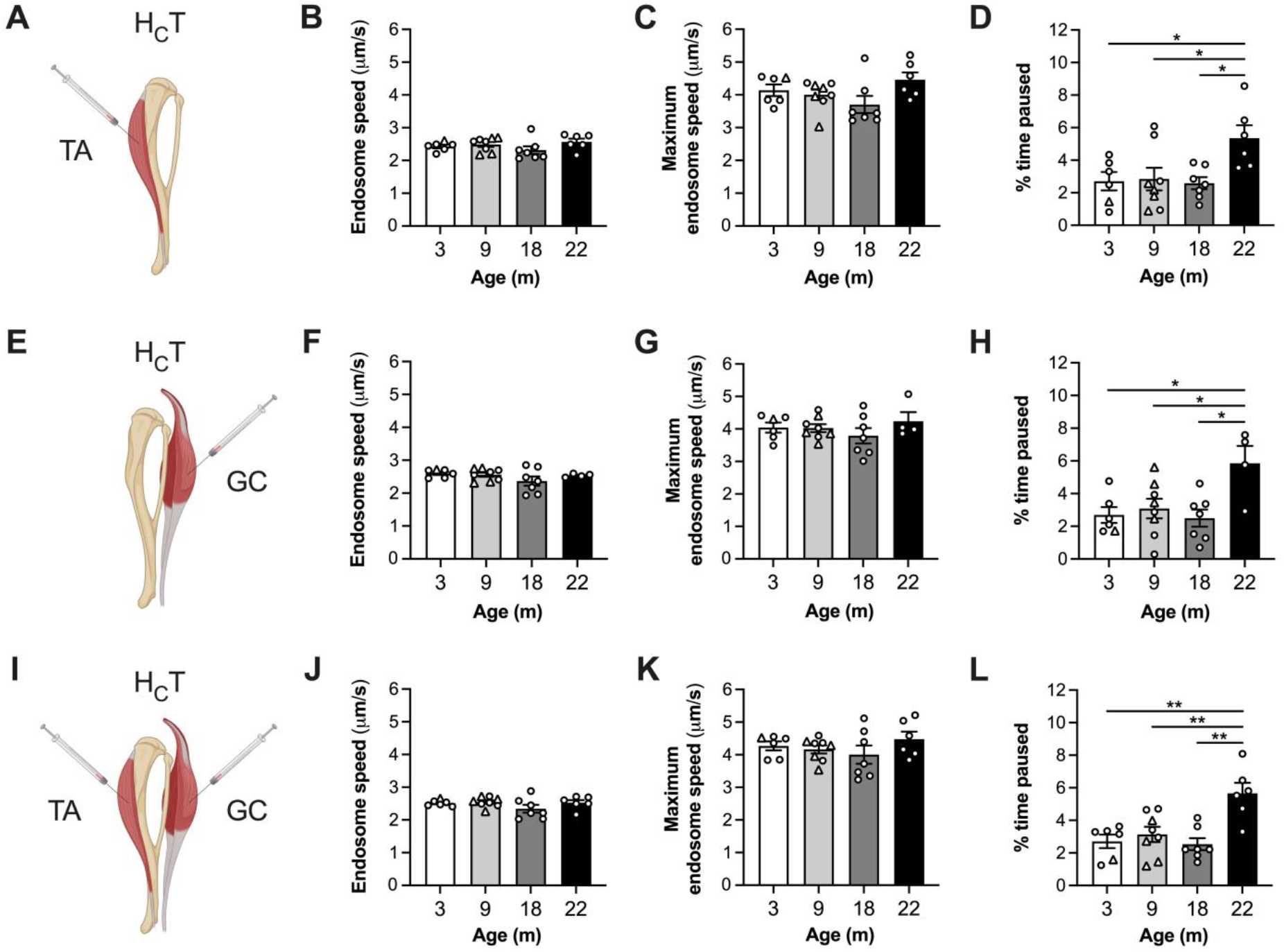
Pausing, but not speed, of signalling endosomes in motor axons is reduced by 22 months of age. (**A-D**) There is no difference in axonal signalling endosome mean (B, *P* = 0.286) or maximum (C, *P* = 0.109 Kruskal-Wallis test) speed within motor neurons innervating the tibialis anterior (TA) from 3 to 22 months of age, but there is an increase in pausing that occurs between 18 and 22 months (D, *P* = 0.018). (**E-H**) There is no difference in axonal signalling endosome mean (F, *P* = 0.245 Welch’s ANOVA) or maximum speed (G, *P* = 0.508) within motor neurons innervating the gastrocnemius (GC) from 3 to 22 months of age; however, an increase in pausing also manifests between 18 and 22 months (H, *P* = 0.014). (**I-L**) Combining the TA and GC data, there is no change with age in signalling endosome mean (J, *P* = 0.252) or maximum (K, *P* = 0.434) speed, but again there is a clear increase in pausing at 22 months (L, *P* < 0.001). *n* = 6-8. All datasets were compared using one-way ANOVA tests, unless otherwise stated. \**P* < 0.05, \*\**P* < 0.01 Tukey’s multiple comparisons test. ○ males; △ females.

### Statistics

Data were assumed to be normally distributed unless evidence to the contrary was provided by the Kolmogorov-Smirnov test for normality. The Bonferroni correction was applied to the multiple Kolmogorov-Smirnov tests performed within each experiment. Bartlett’s test was used to assess equality of variance. Normally distributed data with equal variance between datasets were analysed using unpaired t-tests or one-way analysis of variance (ANOVA) tests followed by Tukey’s multiple comparisons tests, while normally distributed data with unequal variance between datasets were compared using Welch’s ANOVA test followed by Dunnett’s T3 multiple comparisons test. Non-normally distributed data were analysed using a Mann-Whitney U test or Kruskal-Wallis test followed by Dunn’s multiple comparisons tests. All tests were two-sided and an α-level of *P* < 0.05 was used to determine significance. Sample sizes were pre-determined using power calculations and previous experience (Rhymes et al., 2024; Sleigh et al., 2020a; Sleigh et al., 2023), and represent individual animals. Means ± standard error of the mean (SEM) are plotted for all graphs. GraphPad Prism 10 software (version 10.4.0) was used for statistical analyses and figure production.

## Results

Signalling endosomes are specialised organelles that form at axon terminals and retrogradely deliver activated receptors and associated complexes to the cell body to elicit long distance effects, including modulation of gene expression (Villarroel-Campos et al., 2018). We have developed a technique that enables *in vivo* imaging of signalling endosomes as they are transported along polarised microtubules within motor neurons towards the spinal cord (Bilsland et al., 2010; Gibbs et al., 2016; Sleigh et al., 2020b; Tosolini et al., 2021). H_C_T is an atoxic binding fragment of tetanus neurotoxin that can be used to identify BDNF- and TrkB-positive signalling endosomes within axons (Deinhardt et al., 2006). By fluorescently labelling H_C_T and injecting it into mouse hindlimb muscles for uptake at the neuromuscular junction (NMJ), we can subsequently expose the sciatic nerve in anaesthetised mice, and perform time-lapse confocal microscopy to visualise the axonal transport of individual signalling endosomes (**Figure 1A-B**). We prioritise assessment in motor axons over the small percentage of sensory nerves that take up the probe by selecting H_C_T-positive axons with large calibres, which we have previously shown to be motor neurons (Sleigh et al., 2020a).

To determine the impact of aging on axonal transport, we performed intravital imaging in C57BL/6J wild-type mice at 3, 9, 18 and 22 months (**Figure 1C**). The last two timepoints were chosen because many signs that mimic human aging, including loss of muscle mass and function, reduced NMJ integrity, and cognitive decline, begin to manifest between 18 and 22 months in C57BL/6J mice (Borsch et al., 2021; Li et al., 2011; Sheth et al., 2018; Yanai and Endo, 2021), which display an average lifespan of ≈26 and ≈29 months in females and males, respectively (Kunstyr and Leuenberger, 1975).

### Signalling endosomes pause more frequently in aged mice

To begin, we injected H_C_T into the tibialis anterior muscle on one side of the body and into the gastrocnemius on the contralateral side. This enables assessment of signalling endosome axonal transport within motor axons innervating two different muscles, which we have previously shown to have similar baseline transport dynamics (Tosolini et al., 2022), yet display differential vulnerability to trafficking impairment in a mouse model of Charcot-Marie-Tooth disease (CMT) (Rhymes et al., 2024). Hence, the selection of these muscles will enable us to address the possibility that aging may not equally affect the two different populations of motor neurons.

Mice were evaluated at 3, 9, 18 and 22 months, and signalling endosome mean speed, maximum speed and pausing were assessed. Data from the 3 and 9 month timepoints were part of previous studies (Rhymes et al., 2024). We observed no difference in the mean or maximum endosome speeds in motor neurons innervating the tibialis anterior muscle (**Figure 2A-C**). However, there was a significant difference in the percentage time paused, with increased pausing manifesting between 18 and 22 months (**Figure 2D**). Combining all data from 3, 9 and 18 months (since they remain similar) and comparing with 22 months, pausing was increased 1.97-fold at the late timepoint.

Repeating these assessments in motor neurons innervating the gastrocnemius, we again observed no changes in mean and maximum speeds; however, we identified a significant difference between timepoints in the percentage of time paused (**Figure 2E-H**). Comparing all pairwise combinations, we found that significantly more pausing occurred at 22 months than all other timepoints (2.11-fold). It therefore appears that increased endosome pausing (i.e., reduced processivity) with advanced age is common to motor neurons innervating both the tibialis anterior and the gastrocnemius.

Given the similarity in endosome dynamics between the different nerves, we combined the data from the tibialis anterior and gastrocnemius from each animal and re-evaluated transport. Once again, there was no difference in endosome speeds, but pausing was significantly more frequent at 22 months compared to all other time points (2.01-fold, **Figure 2I-L**). Together, these data indicate that the speed with which signalling endosomes are transported within motor axons innervating both the tibialis anterior and the gastrocnemius is resistant to the effects of aging up to 22 months; however, a doubling in the frequency of pausing manifests between 18 and 22 months, which is an early indicator of a decline in processivity of signalling endosome axonal transport during aging.

### Boosting muscle BDNF has no effect on increased pausing at 22 months

BDNF is a secreted neurotrophin that binds and acts through the tropomyosin receptor kinase B (TrkB) receptor, to elicit both local and long-range signalling (Chao, 2003). Once activated, BDNF-TrkB complexes are one of several ligand-receptor pairs that are internalised into signalling endosomes, being retrogradely transported to motor neuron cell bodies, for activating pro-survival gene transcription programmes (Villarroel-Campos et al., 2018). We recently showed that provision of excess BDNF within muscles of CMT mice is able to overcome impaired axonal transport of signalling endosomes caused by dampened BDNF-TrkB signalling (Rhymes et al., 2024; Sleigh et al., 2023). Boosting muscle BDNF can also reduce the baseline level of axonal endosome pausing in wild-type mice (Tosolini et al., 2022), while BDNF-TrkB inhibition in wild-type muscles slows endosome transport and increases pausing (Sleigh et al., 2023). This neurotrophin-receptor pair is therefore required for maintaining physiological signalling endosome delivery to motor neuron cell bodies. There is also evidence that BDNF availability in muscle is reduced with age, at least in rodents (Ming et al., 1999); we therefore hypothesised that intramuscular injections of recombinant BDNF, in a similar manner to previously performed, may reduce the aging-induced increase in endosome pausing.

To test this hypothesis, we compiled data on the dynamics of signalling endosome axonal transport from mice receiving intramuscular injections of H_C_T with BDNF into both the tibialis anterior and gastrocnemius again at 3, 9, 18 and 22 months of age (**Figure 3A**). We chose this paradigm because it provided the most statistically robust pausing defect in the first dataset (**Figure 2D, H and L**). It is worth highlighting that the exposure to increased BDNF is only short-term (4-8 h), but this is sufficient to elicit an acute boost in signalling endosome axonal transport (Rhymes et al., 2024; Sleigh et al., 2023; Tosolini et al., 2022). The data from the timepoints prior to 22 months were generated as part of two previous studies (Sleigh et al., 2020a; Sleigh et al., 2023).

**Figure 3.**
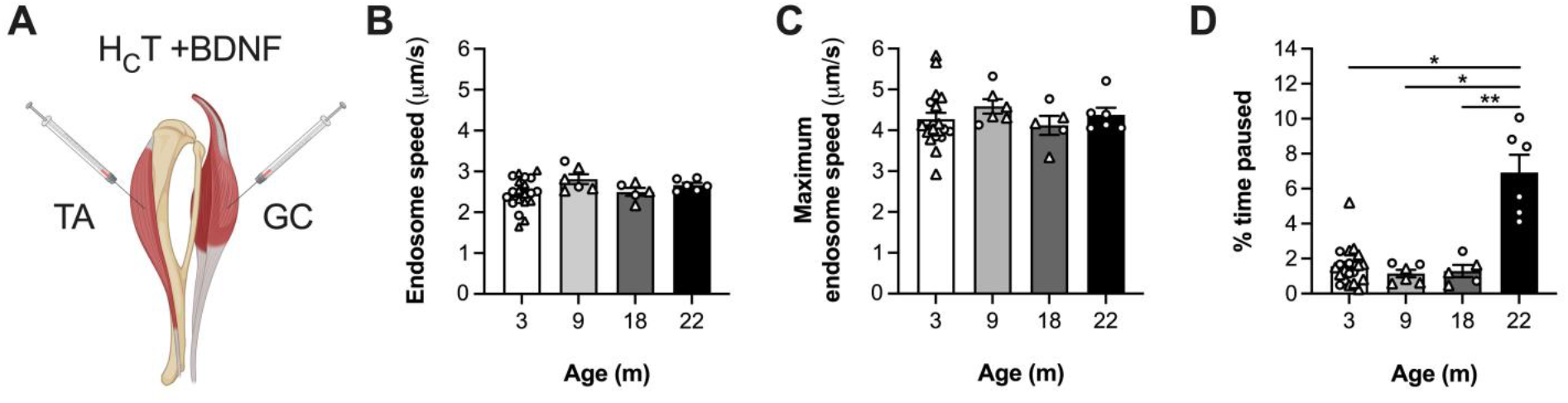
Acute provision of BDNF within aged muscles has no effect on the increased pausing of axonal signalling endosomes. (**A**) Schematic depicting the co-administration of H_C_T and BDNF into both the tibialis anterior (TA) and gastrocnemius (GC). (**B-C**) There is no difference in axonal signalling endosome mean (*P* = 0.124 one-way ANOVA) or maximum (C, *P* = 0.343 Kruskal-Wallis test) speed within motor neurons exposed to increased levels of muscle BDNF from 3 to 22 months of age. (**D**) BDNF is unable to impact the increased pausing in 22 month-old TA- and GC-innervating motor axons (*P* = 0.002 Welch’s ANOVA). *n* = 5-20. \**P* < 0.05 Dunnett’s T3 multiple comparisons test. ○ males; △ females.

Similar to our baseline dataset, we observed no difference in the mean or maximum speed of signalling endosomes across timepoints (**Figure 3B-C**). We again identified a clear difference in the percentage of time that signalling endosomes remained stationary, with greater pausing manifesting between 18 and 22 months (**Figure 3D**), like when excess BDNF was not provided. Compared with all data from 3-18 months, pausing was increased 4.82-fold at 22 months, which is more than twice the increase observed without BDNF injection (**Figure 2L**). This fits with BDNF reducing the amount of pausing pre-22 months and not having an effect at this later age. Indeed, when comparing pausing between untreated mice and those receiving intramuscular BDNF injections, we found decreased pausing upon BDNF administration at 3 and 9 months, with a clear trend towards reduction at 18 months, but not at 22 months (**Figure S1**). Therefore, the age-dependent increase in pausing without affecting endosome speed is replicated in this dataset, suggesting that it is a robust phenotype and one that is impervious to the modifying effects of increased BDNF at aged motor nerve terminals.

## Discussion

Except for very early experiments on the impact of aging on axonal transport performed in cats and dogs (Ochs, 1973) and some later studies in rats (Inestrosa and Alvarez, 1988; Jacob, 1995), it appears there is consensus that cargo trafficking within axons diminishes as animals age; this includes studies in *C. elegans, Drosophila*, mice and rats, and in several types of central and peripheral neurons (Mattedi and Vagnoni, 2019). Nevertheless, work over the last decade in which fluorescent cargoes were individually tracked indicates that the dynamic properties of axonal transport (e.g., velocity, pausing and run length) are not perturbed in old age; rather, it is the frequency of transport that declines. This suggests that the function of the transport machinery is sufficiently preserved over time, but it is the mechanisms that activate and maintain transport, along with the availability of transport components, that are primarily affected by aging.

Here, we add signalling endosomes within intact mouse motor axons to the list of cargoes that are affected by advanced age. Through intravital imaging, we repeatedly observed a two-fold increase in pausing in 22 month-old mice, suggesting that it is a robust and reproducible phenotype. Although the reduced processivity was insufficient to reduce endosome speeds, it is certainly possible that the extra pausing in aged mice reflects the onset of impairments in signalling endosome transport; assessment in older animals (e.g. 24-28 months) will determine whether this is indeed the case. Due to differences in imaging frame rates caused by using two different confocal microscopes (one of them with larger frame sizes, giving more opportunity for an endosome to pause), we were unable to accurately determine the percentage of pausing endosomes across timepoints. However, examining the raw data, we can say that the increase in percentage time paused in older mice was not a result of few endosome pausing many times, but rather more frequent short pausing across the sampled endosomes.

Using a similar imaging approach, we have previously shown that the speed of mitochondrial axonal transport in intact sciatic nerves is also not reduced in mice aged 19-22 months – in fact, for currently unknown reasons, mitochondria were trafficked more quickly in the anterograde direction with a trend towards a retrograde increase (Sleigh et al., 2024). Nonetheless, we did identify that fewer mitochondria were present in aged axons, and that this is possibly a consequence of increased cytoplasmic viscosity in neuronal cell bodies. Unfortunately, due to inherent variability between intramuscular injections and the fact that signalling endosomes are highly processive and do not regularly dock along the axon like mitochondria, it is more challenging to measure the overall flux of signalling endosomes using our imaging approach. Nevertheless, using a radioactive probe similar in function to H_C_T, Lee and colleagues performed computed tomography imaging to show that net retrograde transport of signalling endosomes may indeed progressively decline with age in mice; a clear reduction in probe accumulation within the spinal cord post-injection into the gastrocnemius was identified between 100 and 300 days and again between 700 and 900 days (Lee et al., 2022), mimicking the two distinct phases of decline in NMNAT2-positive vesicle transport reported in aged sciatic, optic and hippocampal nerve explants (Milde et al., 2015). Unfortunately, net retrograde assessments are affected by transport-independent processes (e.g., probe uptake and axon number), thus this approach may therefore overestimate the decline in trafficking with age, as suggested by the lack of change in signalling endosome dynamics observed from 3 to 18 months (90 to 615 days). That being said, our data are consistent with observations across models that there is a general decline in the total amount of axonal transport. To provide additional data to support this, in the future we could perform sciatic nerve ligation prior to intramuscular H_C_T injections, and assess the amount of probe accumulation distal to the ligation, as has been done previously with tetanus neurotoxin fragments (Roux et al., 2006). Nonetheless, this approach would also be impacted by the decline in NMJ integrity with age (Li et al., 2011; Valdez et al., 2010), resulting in reduced H_C_T uptake. This raises the important point that age-dependent decline of the neuromuscular system may also in part contribute to why we do not see a clear disruption in axonal transport speeds with age, since H_C_T will not accumulate and be trafficked into declining axons that have lost connection to the muscle.

The reason for the enhanced pausing of signalling endosomes at 22 months is currently unknown. The increased cytoplasmic viscosity we identified in cell bodies of aged peripheral nerves is unlikely to directly impact the retrograde trafficking of endosomes since they are formed in the periphery (Sleigh et al., 2024). However, signalling endosomes and mitochondria frequently interact within axons (Cioni et al., 2019), so there is the possibility for a transfer of signals from anterogradely-delivered mitochondria. Moreover, although there was no clear increase in viscosity of the axoplasm of aged peripheral nerves, we did detect a small decrease in particle diffusiveness (Sleigh et al., 2024), suggesting that spatial hinderance/local crowding in distal axons (e.g., from aggregated proteins known to accumulate in aging (Trigo et al., 2019)) may partially contribute to the rise in pausing. Supporting this idea, experimentally reducing the availability of axonal mitochondria causes an acceleration in protein accumulation within aged axons (Vagnoni et al., 2016).

The BDNF receptor, TrkB, is downregulated at the mouse NMJ by 24 months of age (Personius and Parker, 2013) and the enhancing effect of BDNF on neuromuscular transmission is lost at 24, but not 18 months of age (Greising et al., 2015a). Furthermore, reduced TrkB function results in impaired NMJ structure, neurotransmission, and muscle function, mimicking the impact of aging on the neuromuscular system (Gonzalez et al., 1999; Greising et al., 2015b; Kulakowski et al., 2011). Together, these findings suggest that an age-related decline in BDNF-TrkB signalling, which is known to regulate endosome transport speeds within motor neurons *in vivo* (Sleigh et al., 2023), may have a role in the identified pausing phenotype. Consistent with this, intramuscular injection of BDNF reduced baseline pausing at 3, 9 and 18 months (with the latter being only a trend), similar to previous observations (Tosolini et al., 2022); however, BDNF had no impact on pausing when administered at 22 months, similarly to that observed in mice modelling amyotrophic lateral sclerosis (Tosolini et al., 2022). This suggests that decreased TrkB availability at aged motor nerve terminals may account for the increased pausing at baseline and provides an explanation for why acutely boosting muscle BDNF does not impact pausing at 22 months. This also fits with the finding that in human motor neurons, BDNF controls transcriptional networks that mediate cytoskeletal organisation and axonal regeneration (Vargas et al., 2023), which are two processes perturbed by aging (Geoffroy et al., 2016; Richardson et al., 2024; Verdu et al., 2000). Furthermore, BDNF has been shown to both increase local translation of cytoplasmic dynein (Villarin et al., 2016) and enhance its recruitment to signalling endosomes (Mitchell et al., 2012), deficits in which may directly contribute to the reduced processivity of signalling endosomes in aged axons through reduced engagement of dynein motors. Intriguingly, demyelination causes an increase in speed of anterogradely transported mitochondria as a means of neuroprotection (Licht-Mayer et al., 2020), which mimics what we observed in aged peripheral nerves *in vivo* (Sleigh et al., 2024); Schwann cells are a source of BDNF, suggesting that this neurotrophin could also play a role in the mitochondrial phenotype we previously identified.

In summary, we show that the processivity of signalling endosomes is consistently reduced in motor axons of aged mice, manifesting as a doubling in the amount of pausing between 18 and 22 months. This may result from a decline in TrkB availability at the NMJ, which is consistent with the observed loss of response to increased muscle BDNF in the aged motor nervous system.

## Acknowledgements

We thank the personnel of the Denny Brown Laboratories (UCL Queen Square Institute of Neurology) for assistance in maintaining the mouse colonies. This project was funded by Medical Research Council awards (MR/S006990/1 and MR/Y010949/1) (JNS); a Wellcome Trust Sir Henry Wellcome Postdoctoral Fellowship (103191/Z/13/Z) (JNS); a Rosetrees Trust grant M806 (JNS, GS); the UCL Neurogenetic Therapies Programme funded by The Sigrid Rausing Trust (JNS); the UCL Therapeutic Acceleration Support scheme supported by funding from MRC IAA 2021 UCL MR/X502984/1 (JNS); a Junior Non-Clinical Fellowship from the Motor Neuron Disease Association (Tosolini/Oct20/973-799) (APT); AFM-Téléthon (BM), Kennedy’s Disease Association (BM); Brain Research Trust (BM); Kennedy’s Disease UK (BM); UCL Queen Square Institute of Neurology Kennedy’s Disease Research Fund (BM); the Horizon 2020 Framework Programme (Grant Agreement No. 857524) (AV); Wellcome Trust Senior Investigator Awards (107116/Z/15/Z and 223022/Z/21/Z) (GS); and a UK Dementia Research Institute award (UKDRI-1005) (GS). The graphical abstract and Figures 1A, 1C, 2A, 2E, 2I and 3A were created using https://www.biorender.com.

## Author Contributions

Conceptualisation: AV, GS, JNS. Investigation: DV-C, ERR, APT, JNS. Writing – Original Draft: JNS. Writing – Review & Editing: DV-C, ERR, APT, BM, AV, GS, JNS. Resources: BM. Supervision: GS, JNS.

## Conflict of Interest

None declared.

## Supplementary Information

### Supplementary Tables

**Table S1.** Experimental sample sizes and mouse details. *BDNF*, brain-derived neurotrophic factor; *d*, days; *F*, female; *M*, male; *m*, month; *n*, sample size.

| Time point (m) | BDNF (-/+) | *n* | Sex | Age (d) | Reference |
| --- | --- | --- | --- | --- | --- |
| 3 | - | 6 | 5M/1F | 91-96 | (Rhymes et al., 2024) |
|  | + | 20 | 7M/13F | 90-115 | (Sleigh et al., 2020; Sleigh et al., 2023) |
| 9 | - | 8 | 4M/4F | 271-279 | (Rhymes et al., 2024) |
|  | + | 6 | 2M/4F | 251-271 | (Sleigh et al., 2020) |
| 18 | - | 7 | 7M/0F | 555-593 | This study |
|  | + | 5 | 2M/3F | 588-615 | (Sleigh et al., 2020) |
| 22 | - | 6 | 6M/0F | 663-687 | This study |
|  | + | 6 | 6M/0F | 664-678 | This study |

### Supplementary Figures

**Figure S1.**
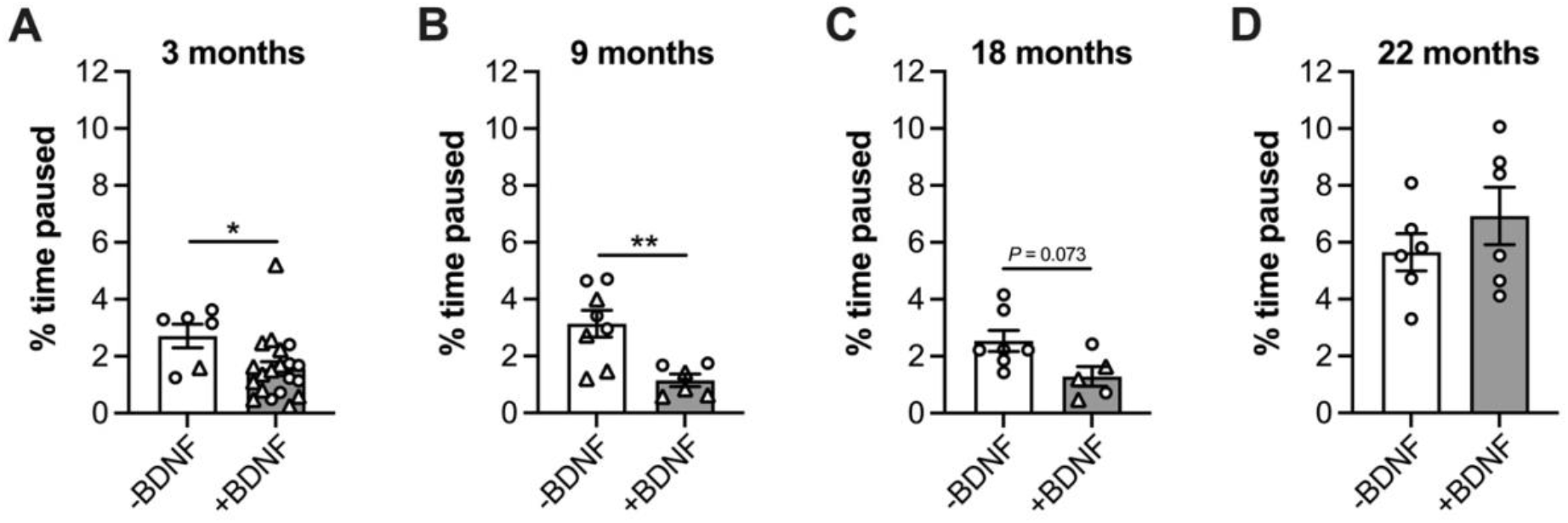
The modifying effect of boosting muscle BDNF on axonal endosome pausing is lost at 22 months. Intramuscular injection of BDNF reduces pausing in 3 (\**P* = 0.033 unpaired *t*-test) and 9 (\*\**P* = 0.005 unpaired *t*-test) month-old mice, and causes a trend towards less pausing at 18 months (*P* = 0.073 Mann-Whitney *U* test); however, there is no such decrease at 22 months (*P* = 0.315 unpaired *t*-test). *n* = 5-20. ○ males; △ females.

## Notes

### Competing Interest Statement

The authors have declared no competing interest.

